# Plants perceive aerosols as an intensification of atmospheric dryness and react according to their isohydricity

**DOI:** 10.1101/2023.12.15.571659

**Authors:** Chia-Ju Ellen Chi, David A. Grantz, Juergen Burkhardt

## Abstract

Hygroscopic aerosols deposited to leaves are a local water vapor sink and can affect the water balance of plants by deliquescence and the formation of hydraulic films that penetrate into the stomata. Stomatal responses to aerosols and vapor pressure deficit(VPD) were investigated in two poplar clones grown hydroponically in ventilated greenhouses with and almost without ambient aerosols.

With increasing VPD, transpiration increased in ANI, the more anisohydric clone, and decreased in ISO, the more isohydric clone, while aerosols had little effect. In ANI, stomatal conductance (g_sw_) and photosynthesis (A) decreased slightly with increasing VPD, but significantly with exposure to aerosols. Leaf carbon isotopes confirmed the long-term reduction in stomatal aperture by aerosols. In ISO, g_sw_ and A decreased strongly with increasing VPD. Aerosols had no effect on stomatal conductance in ISO, but increased the minimum leaf conductance and decreased the turgor loss point. In both clones, aerosols reduced stomatal density by >20%, indicating increased water scarcity.

Aerosols enhance the transmission of atmospheric dryness to the leaf, with plant responses depending on their isohydricity. Sensitive stomatal closure of isohydric plants is an effective adaptation to atmospheric dryness, but aerosol accumulation mediates a liquid pathway for water loss that undermines stomatal control.

## Introduction

Theory and models of plant gas exchange regard leaf surfaces as clean surfaces that do not interact with the gases flowing past them. In reality, however, there are deposited aerosols on every leaf surface, a large proportion of which are hygroscopic. This experimental study investigates the extent to which deposited ambient aerosol affects the gas exchange and physiological parameters of two hydroponically grown poplar clones. The expression “aerosols” in this paper includes deposited aerosols, although they are no longer in the airborne state.

### Hygroscopic aerosols

Atmospheric aerosols are a natural part of the environment, ranging from few nanometers to about 100 μm. Particles across this size range are found on foliar surfaces. Deposition to vegetation is a function of aerodynamic diameter and canopy surface characteristics (Lindberg *et al*., 1986). Aerosol comprises a dynamic mixture of solid and liquid particles, originating from natural sources such as sea salt, geologic dust, and volcanic ash; as well as anthropogenic sources including combustion, construction and agricultural production (Hamilton, 2015; Burkhardt & Grantz, 2017; Lu & Tian, 2017; Burkhardt *et al*., 2018; Chappelka & Neufeld, 2018). Aerosol loading of the troposphere and deposition to terrestrial vegetation are increasing along with the hygroscopicity of ambient aerosol in the Anthropocene (Andreae, 2007), which has been shown to enhance crop canopy photosynthesis (Chameides *et al*., 1999) and tree stem growth rates, associated with increased diffuse radiation, lower VPD, and reduced midday leaf temperatures (Steiner & Chameides, 2005; Wang *et al*., 2018; Wang *et al*., 2021).

The majority of aerosols in the atmosphere are hygroscopic and function as cloud condensation nuclei (CCN) and, e.g., more than 30% of the European atmospheric aerosol mass consists of inorganic ions (mainly ammonium, sulfate, nitrate; Bressi *et al*., 2021). These particles do not lose their hygroscopic properties when depositing to plant surfaces, and particularly salts will deliquesce within the humid leaf boundary layer, many of them at 70% to 80% relative humidity (RH). Such values are often reached on leaf surfaces, particularly close to transpiring stomata. Saturated or highly concentrated solutions develop and form thin films with minute amounts of liquid water that can be present on leaves even during hot, dry summer days (Burkhardt and Eiden, 1994; Burkhardt and Hunsche, 2013). One important property of saturated solutions is that the saturation humidity in the immediately adjacent air is equal to the deliquescence humidity of the salt at the respective leaf temperature (e.g., 75% RH over NaCl; Winston and Bates, 1960). Another important property of salts is their immediate response to changes in relative humidity (Wylie, 1955) through absorption and desorption of water.

### Hydraulic activation of stomata (HAS) and its detection

A liquid transport pathway into the stomata may develop, which is derived from the thin aqueous film that coats all surfaces in the presence of atmospheric water vapor (Wylie, 1955; Ewing, 2005). In the presence of deliquescent salts, here derived from ambient aerosol deposited to foliar surfaces (Burkhardt & Grantz, 2017), additional solution is added to this film. Repeated drying and rehydration of the saturated aqueous films lead to salt creep (Qazi *et al*., 2019), and on the otherwise hydrophobic waxy cuticle this process is particularly effective for chaotropic ions that reduce the surface tension of water (Burkhardt et al., 2012). As a result the solutions spread across the leaf surface, as is visible in electron micrographs, eventually entering into the pores along the stomatal walls (Burkhardt & Pariyar, 2014; Grantz *et al*., 2018), (see videos in Burkhardt & Pariyar, 2014) a linkage that has been demonstrated repeatedly for foliar fertilization, herbicide uptake, and nanoparticle infiltration (Eichert *et al*., 1998; Eichert & Goldbach, 2008; Basi *et al*., 2014). Through this process of hydraulic activation of stomata (HAS; Burkhardt, 2010), a thin film is provided that can also serve as a pathway for liquid water flow from leaf interior towards the leaf surface, where it evaporates and is replenished from apoplastic water within the leaf, acting similar to a wick. The water flux by this pathway is of unknown magnitude but may be large and is not reduced by stomatal closure, which functions effectively to reduce diffusional conductance to water vapor.

Recently, greenhouse sap flow measurements of hydroponic sunflowers showed that ambient aerosols caused almost 50% higher transpiration rates (E) when comparing plants grown in almost aerosol free air with plants grown in unchanged air (Burkhardt et al., 2023). Plant conductance G was twice as high at low VPD and the difference disappeared with increasing VPD, supporting the wick action of HAS and adjacent leaf surface areas that (at low VPD) contributed to evaporation of liquid water from the leaf surfaces. Aerosols thus caused effects which were similar to effects of VPD, and plants grown in the ambient air (AA) greenhouse reached the overall mean transpiration rates observed at 1.68 kPa in filtered air (FA) at the lower VPD of 0.34 kPa. In spite of hydroponic cultivation with unlimited water supply, sunflowers grown in ambient air developed several indicators of water scarcity when compared to plants grown in filtered air: lower osmotic potential, limitation of triose phosphate utilization (TPU), higher leaf mass per area (LMA) and distinct tracer transport patterns in the leaf (Burkhardt et al., 2023).

Although the detailed processes of wick action have not been elaborated yet (necessary conditions would include a continuous thin film and appropriate distributions of the water potentials of apoplast, salt solutions, and the atmosphere), it is clear that the direction of flux happens always along the water potential gradient. Short-term fluctuations of wind, temperature, and humidity will contribute to reduction in water potential near the salt solution and and Marangoni forces may be involved through changes in surface tension (Shao et al., 2022). The leaf surface surrounding stomata would in this case participate in transpiration by sorption and desorption of water. This would create a transpiration differential compared to aerosol-free leaves, and potentially a bi-phasic response as the effect diminishes at high VPD.

Gas exchange measurements rather than sap flow are used in the present study for the detection of HAS. These are not trivial measurements because the incremental flux along thin aqueous films appears in the total water flux and cannot be easily isolated. The most direct experimental proof is the simultaneous measurement of gas exchange and stomatal pore aperture, which requires sophisticated equipment and the evaluation of several thousand stomatal images (e.g., Burkhardt et al., 2001; Grantz et al., 2018). Whereas theory predicted the determination of stomatal conductance by stomatal aperture, in *Vicia faba* leaves grown in ambient or in filtered air proved that aerosol decreased stomatal apertures at each level of VPD and increased stomatal conductance at each level of aperture (Grantz et al., 2018), and aerosols reduced the magnitude of variability between pores (“patchiness”; Grantz et al., 2020).

Under certain conditions, foliar carbon isotope composition might be used for HAS detection. Photosynthetic carbon isotope discrimination from leaves can be interpreted as integrated gsw under which carbon fixation occurs, if other conditions are equal (Farquhar & Richards, 1984). Less negative δ^13^C indicates reduced stomatal opening, typically associated with water deficit, reduced gsw, and increased WUE (Marron et al., 2005; Monclus et al., 2006; Broeckx et al., 2014).

The Ball-Berry (BB) approach (Ball et al., 1987), could be an alternative possibility for HAS detection, because it integrates of both transpiration and photosynthetic fluxes in one empirically derived formula. The empirical BB approach describes how g_sw_ is determined on the basis of its apparent linear sensitivity to a lumped environmentally weighted term for photosynthesis that includes relative humidity (Medlyn et al., 2011). This has been very successful in predicting gsw, but has been criticized for its empirism and lack of theoretical basis for the proportionality of gs/A and relative humidity (e.g., Monteith, 1995; Medlyn et al., 2011). However, in combination with the HAS theory, the relative humidity at the leaf surface (Hs) may become apparent (Burkhardt, 2010).

### Possible consequences of HAS for models of gas exchange

Salts react directly to relative humidity by attracting a thermodynamically defined amount of liquid water, which can be measured by electrical conductivity – a classical method to determine relative humidity (Wylie, 1955). As relative humidity thus translates to a defined amount of liquid water, there is no contradiction to the Helox experiments that seemed to show that stomata react to transpiration and not relative humidity (Mott and Parkhurst, 1991). At the same time, a liquid water connection across the stomata provides a functional humidity sensor, where leaf constituents inside the stomata get direct hydraulic signals, with information about the atmospheric humidity state outside of the stomata and independent of the actual stomatal aperture. This enables the so-called “feedforward” effect, in which plants not only decrease conductance but also transpiration in response to increasing VPD (Farqhuar, 1978). This phenomenon has long been enigmatic (Schulze et al. 1972, Farquhar 1978, Maier-Maercker 1983, Grantz 1990, Mott and Parkhurst 1991,

Monteith 1995, Franks et al. 1997, Buckley 2019). Decreasing transpiration is expected to lead to an increase in water potential and plants would require an external humidity sensor or stomatal patchiness (spatially heterogeneous stomatal closure) to further decrease stomatal conductance (Mott and Peak 2007). HAS and the connected water loss by wicking can provide a mechanistic explanation. Deliquescence of deposited aerosols may serve as a humidity sensor (Burkhardt, 2010), and aerosol exposure may influence patchiness, as shown in *V. faba* (Grantz *et al*., 2020).

With an established liquid water connection across open stomata, a second pathway of transpiration parallels the diffusional vapour flux. Stomatal aperture only controls outgoing water vapour flux and incoming CO_2_ flux, but not the additional transpiration along the liquid pathway. This creates a bias in those models of gas exchange, which rely on the equivalence of the stomatal passage by diffusion, scaled only by the ratio of molecular diffusivities in air usually taken as 1.6 (Ball et al., 1987; Damour et al., 2010; Medlyn et al., 2011).

The thin films along the stomatal throat do not disappear and can still be active in liquid water transport when stomata close. They could be the main mechanistic reason for “leaky” or “malfunctioning” stomata that have often been described (Kerstiens, 1996; Duursma et al., 2019) and represent the minimum leaf conductance (g_min_) measured in darkness. Increasing g_min_ caused by hygroscopic leaf surface material has repeatedly been documented for a number of plant species, for ambient aerosols and salt sprays (Burkhardt & Pariyar, 2014; Burkhardt *et al*., 2018; Grantz *et al*., 2018; Chi *et al*., 2022). The increase in g_min_ is evidence that the liquid transpiration along the established film from HAS is still functional under these conditions. Any such leakage, uncontrolled by stomatal closure, may become a challenge for plants, because the water potential gradient is extremely steep between the substomatal cavity (typically -7 MPa, 95% relative humidity) and the lowest part of the atmosphere, just outside the leaf (−70 MPa, 60% rh). Such leakage is a critical factor for tree mortality and is poorly quantified (Brodribb et al., 2020).

### Isohydricity

Plants regulate diffusional loss of water vapor, the dominant component of transpiration (E) along a continuum of isohydricity, implying regulation that either favors prolonged stomatal opening, thus sustaining carbon acquisition and leaf cooling, or sensitive stomatal response to rising VPD or drying soil, thereby maintaining leaf water status (Mitchell *et al*., 2013). Isohydric plants exhibit strict stomatal regulation and large declines in stomatal conductance (g_sw_) and net photosynthesis (A) but minimal changes in leaf water potential (Ψ), during midday and drought. Anisohydric plants maintain g_sw_ and A across a wide range of Ψ at the risk of photosynthetic inhibition and enhanced vulnerability to hydraulic failure (Klein, 2014; Meinzer *et al*., 2016; Meinzer *et al*., 2017).

While the wick hypothesis worked well for sunflowers which are strongly anisohydric (Tardieu and Simmoneau, 1998; Burkhardt et al., 2023), there were indications that not all plants would transpire more in response to aerosol exposure. Burkhardt and Pariyar (2016) found that the anisohydric species beech showed linearly increasing sap flow with increasing VPD, with slopes doubling for ammonium sulfate-sprayed and tripling for sodium nitrate-sprayed beech compared to control seedlings, whereas the isohydric species Scots pine showed constant transpiration rates with increasing VPD, independent of the aerosol.

HAS enhancement of water loss and the complexity of isohydricity has received limited attention (Burkhardt & Pariyar, 2016), and is tested here by an aerosol exclusion experiment with hydroponically grown poplars in two parallel greenhouses, one receiving unfiltered, ambient air (AA), the other one receiving filtered air (FA) almost without aerosols. Here we use different clones of hydroponically grown poplar to broaden our knowledge on the spectrum of plant responses to ambient aerosols.

### Relevance of aerosols and HAS in tree mortality

If hygroscopic aerosols act as a water vapor sink and HAS transfers atmospheric dryness to the leaf interior, plant exposure to VPD increases. An important implication is that the effect of increasing VPD on plants (Novick et al., 2016, Grossiord et al., 2020) is amplified. In this case climate change and air pollution could reinforce each other in their effects on regional forest dieback/tree mortality.

Large regional forest diebacks occurred mainly in the 1980s and again over the last 20 years, now mostly named tree mortality. Forest decline in the 1980s was considered to be caused by air pollution, which led to high acidity and nutrient inputs into forests and resulted in nutrient imbalances (Schulze, 1989). Due to their small mass, the deposition of aerosols contributed only little to overall input, while their hygroscopic action was neglected. Although typical damage patterns already resembled drought at that time, a mechanism for how desiccation could be caused by air pollution was not understood.

The large-scale regional tree mortality observed throughout the last 20 years has been attributed to climate change, and a consensus is emerging that particularly accumulated plant exposure to rising VPD plays a crucial role (Breshears et al., 2005; Allen et al., 2010; McDowell et al., 2022). Air pollution is almost neglected in this discussion, although global concentrations of aerosols and ammonia, an important precursor gas, have been constantly rising throughout the last decades, even though emissions in some regions started to decrease (Crippa et al., 2018).

Aerosols are globally distributed and their interaction with plants via HAS is a mostly physicochemical effect, which would support a possible contribution to global tree mortality. They can contribute by causing higher transpiration rates, or could act by the increase in minimum conductance, which has meanwhile been observed for a number of species. They might thus also have a hidden impact on isohydric plants, which have a greater tendency to suffer from tree mortality than anisohydric plants (McDowell et al., 2008).

### Hypotheses

Hypothesis 1: Aerosols at ambient air concentration intensify the transmission of atmospheric dryness to the leaf interior.

Hypothesis 2: The plant response to hygroscopic aerosols depends on isohydricity, as determined from their responses to drought.

Hypothesis 3: Aerosols are the mechanistic reason for the proportionality of gs/A and relative humidity as considered in the Ball-Berry approach, and for the enigmatic feedforward effect in stomatal response to VPD.

Hypothesis 4: The impact of aerosols to isohydric plants can be more substantial than to anisohydric plants.

### Experimental approach

Whereas the physical conditions for the HAS hypothesis have been relatively well elaborated, the actual relevance for plants is still unclear but seems to depend on isohydricity (Burkhardt & Pariyar, 2016). While expecting differential responses from plants with varying isohydricity, the degree of isohydricity was not our main experimental focus. We chose poplar clones to avoid genetic variability and because poplars are known to cover a considerable part of the isohydricity spectrum (Attia et al., 2015). Plants were hydroponically grown in order to concentrate on the aerial part of plant water relations.

We used the two identified poplar clones to investigate short-term (gas exchange, water potential) and long-term (g_min_, δ^13^C, TLP, stomatal density/leaf surface morphology) responses to increasing VPD and aerosols. For evaluating the VPD influence we used VPD curves from 0.75 to 1.75 kPa expanding upon methods and results of Chi *et al*. (2022). For ambient aerosols, we compared plants grown in parallel ventilated greenhouses with and without aerosol exclusion by filtration. Water use efficiency (WUE) and the descriptive parameters of the Ball-Berry model (Ball *et al*., 1987) were obtained and compared with direct measurements of related parameters (Medlyn *et al*., 2017; Yi *et al*., 2019; Yuan *et al*., 2019; Pieters *et al*., 2022). Scanning electron microscopy (SEM) images were obtained to evaluate leaf surface characteristics and stomatal structures. From the evaluation we found that aerosols consistently were perceived by the plants as additional atmospheric dryness and created water scarcity in both clones, despite hydroponic cultivation. The clones developed contrasting symptoms, depending on the isohydricity of the clone.

## Materials and Methods

### Plant material and growth conditions

Two hybrid poplar clones (isohydric clone: *Populus maximowiczii x nigra* and anisohydric clone: *Populus trichocarpa x maximowiczii*), which appeared to differ in isohydricity (Xu et al., 2018), were grown in hydroponic solution containing all essential nutrients. Solutions were renewed weekly to avoid nutrient depletion in the root zone. The clones differed in isohydricity with *P. maximowiczii x nigra* (clone ISO) exhibiting greater isohydric behavior and *P. trichocarpa x maximowiczii* (clone ANI) more anisohydric responses.

Poplar seedlings were assigned randomly to one of two greenhouses, one ventilated with ambient air (treatment AA, n = 12), and the other with filtered air (treatment FA, n = 12). Filtration in this greenhouse system removed about 99% of ambient aerosols (Grantz *et al*., 2018). The relative humidity and temperature of the greenhouses were recorded as described previously (Pariyar *et al*., 2013; Burkhardt & Pariyar, 2016). Temperature and leaf to air vapor pressure deficit (VPD) in the FA greenhouse were higher and relative humidity was lower than in the AA greenhouse, due to flow restrictions associated with filtration. Average temperature in AA was 24.49 ± 0.18°C, R.H. 60.23 ± 0.50%, and VPD 1.72 ± 0.04 kPa; and in FA 25.89 ± 0.16°C, 48.47 ± 0.36%, and 2.16 ± 0.04, respectively (Figure 1).

**Figure 1.**
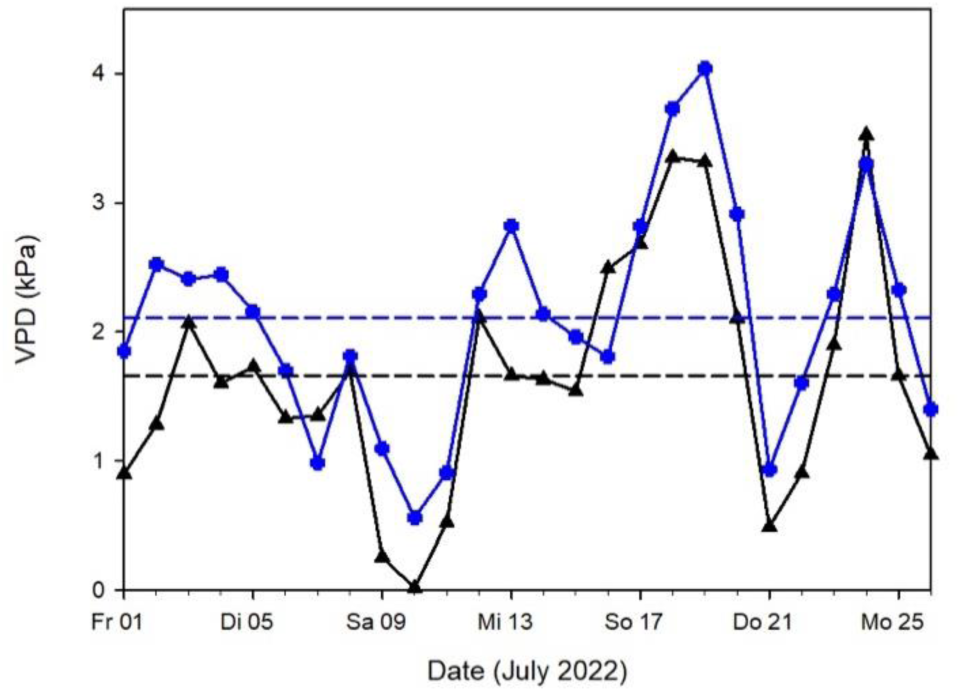
Daily values and means of VPD in the greenhouses throughout the 2022 measurements. Black lines: AA, blue lines: FA; dashed lines: mean values.

All measurements were obtained July to August 2021 and 2022, when plants had branched and attained a height of about 160 cm. All sampling was conducted on the youngest fully-expanded leaf of each branch.

### Gas exchange measurements

The responses of stomatal conductance for water vapor (g_sw_) to increasing vapor pressure deficit (VPD) at constant leaf temperature (25°C) were determined using a Portable Photosynthesis System (LI-6800; LICOR Biosciences, Lincoln, NE, USA). For these response curves, photosynthetic photon flux density (PPFD) was set to 1000 μmol m^−2^ s^−1^ to avoid photoinhibition, and sample cell CO_2_ concentration was held constant at 400 μmol mol^−1^. Gas exchange parameters were measured as a stepwise sequence of VPDs, as follows: 0.75 kPa (2 hr), 1.00 (1 hr), 1.25 (1 hr), 1.50 (1 hr), 1.75 (1 hr) kPa, allowing time for steady state conditions to be established.

The response curves to increasing VPD represented means of stomatal conductance for water vapour (g_sw_), transpiration rate (E), and net photosynthesis rate (A) at each VPD (n = 3). In further analysis, g_sw_ was normalized to g_sw_ at the lowest VPD (0.75 kPa) and as the ratio of aerosol treatments at each VPD.

Intrinsic water use efficiency (WUE) (n = 3) was calculated as in Equation (1), at each VPD level: 0.75, 1.00, 1.25, 1.50, and 1.75 kPa, respectively (Medlyn *et al*., 2017).

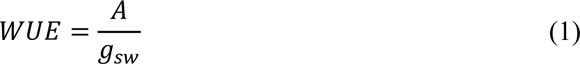

The Ball-Berry model of gas exchange parameters (Equation (2)) was run using measured g_sw_ (n = 3) from the VPD curves.

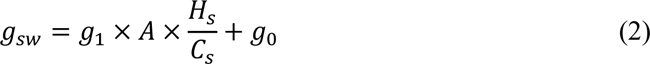

This model considers g_sw_ as a function of A, relative humidity at the leaf surface (H_s_), and CO_2_ concentration at the leaf surface (C_s_, μmol mol^−1^). The slope (g_1_) and intercept (g_0_) between measured g_sw_ and 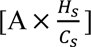 (the Ball-Berry Index) were calculated. The g_1_ represents a compromise between the costs and benefits of increasing g_sw_ relative to the photosynthetic activity of the leaf. The intercept, g_0_, represents an estimate of minimum leaf conductance (mol m^−2^ s^−1^) (Ball *et al*., 1987; Franks *et al*., 2017; Medlyn *et al*., 2017; Miner & Bauerle, 2017). These estimates were compared with directly measured leaf minimum conductance (g_min_) and WUE.

To avoid artifact due to endogenous diurnal rhythms (McClung, 2006; Nozue & Maloof, 2006), all gas exchange measurements were conducted during similar hours of the day, and alternated between AA and FA.

### Cuticular and stomatal characteristics

Leaf surface characteristics were visualized by scanning electron microscopy (SEM, Leo 1450VP, Zeiss, Jena, Germany). Fresh leaves were obtained from the two clones in both greenhouses, sealed in plastic zipper bags, and transferred immediately to the laboratory. Samples were sealed with laboratory glue at the incisions to prevent water evaporation, and the surface was coated with palladium for high-vacuum SEM imaging. Images were processed using ImageJ (Schneider *et al*., 2012).

The adaxial leaf surfaces contained few stomata. Abaxial leaf surfaces were analyzed for the length, width, and area of the stomatal complexes, and for stomatal density (n = 10). In order to test if aerosols had an influence on leaf surface hydrophobicity, contact angles were determined. 1-μl droplets of water were placed on the cuticles and contact angles were measured using a goniometer (DSA 30E; Kruess GmbH, Hamburg, Germany) (n = 18). The surface tension of the water and the contact angles were determined by the pendant drop method (Burkhardt *et al*., 2012).

### Carbon isotope discrimination

The carbon isotope composition was measured with an isotope ratio mass spectrometer (IRMS, C-N-S Analyzer, and MS-2020; SerCon Ltd., Crewe, UK). The harvested fresh leaves were dried to constant weight and ground to fine powder (n = 8). 1 mg of ground sample was loaded into a tin capsule and subjected to oxidation. The purified gas stream was fractionated by gas chromatography and the CO_2_ was brought to the mass spectrometer. The ^13^C value was calculated, and carbon isotope discrimination ( 𝛿 ^13^C) determined by comparison to a standard (Condon *et al*., 2002; Burkhardt & Pariyar, 2016).

### 4.1. Leaf water potential at noon and at turgor loss point

Total leaf water potential was obtained at noon (Ψ_noon_, MPa). Fresh leaves were wrapped in aluminum foil then excised from both clones in both greenhouses. Samples were obtained in succession immediately prior to measurement. Scholander Pressure Chamber (Model 3005, Soil Moisture Equipment Corp., Santa Barbara, CA, USA) (n = 5) was used as previously described (Pariyar *et al*., 2013; Chi *et al*., 2022).

Leaf water potential at the turgor loss point (𝜋_tlp_, MPa) was determined using the method of Bartlett *et al*. (2012a). Leaf discs were prepared from excised fresh leaves with a 6 mm diameter cork borer. Discs were quick-frozen in liquid nitrogen to fracture the cell walls (n = 8) and punctured to disrupt the cuticle, and then rapidly sealed in a vapor pressure osmometer (VAPRO 5600, Wescor, Inc, Logan, UT, USA). The osmolality C_o_ (mmol kg^−1^) was recorded at equilibrium, and converted to osmotic potential (𝜋_o_, MPa) using the Van’t Hoff Equation (3). C_o_ is the molar solute concentration (mmol kg^−1^), R is the universal gas constant (m^3^ MPa K^−1^ mol^−1^), and T is temperature (K) (Khare, 2015). Based on the strong correlation between 𝜋 _o_ and 𝜋 _tlp_ (R^2^ = 0.91), 𝜋 _tlp_ was calculated using Equation (4) (Bartlett *et al*., 2012b; Sjoman *et al*., 2015; Banks & Hirons, 2019).

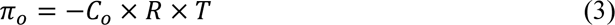

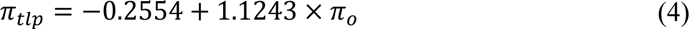

### Minimum leaf conductance

Fresh leaves were excised from both clones in both greenhouses. The petioles were sealed with molten paraffin. Leaves were held in darkness and weighed every 90 minutes using a digital balance (EX125M, EXPLORER TSEMIMICRO, Ohaus Corporation, Parsippany, NJ, USA; 120 g x 0.01 mg). Temperature and humidity were continuously recorded to determine VPD and leaf area was analyzed with ImageJ. Minimum leaf conductance (g_min_, mmol m^−2^ s^−1^) was calculated as described (Sack & Scoffoni, 2011; Duursma *et al*., 2019).

### Data processing and statistical analysis

Based on preliminary research (Chi *et al*., 2022), the minimum sample size to achieve an alpha of 0.05 and power of 80% of each measurement was estimated by using the *pwr* package in R Studio (R version 4.0.3). The sample size then resulted in the range from n = 3 to 8. The Shapiro–Wilk test was used to test data for normality. The F-test or Fligner– Killeen test was performed to compare the homogeneity of variances. For normally distributed data, mean separation was evaluated by two-tailed Student’s t-test. For non-normally distributed data, the difference between medians was evaluated nonparametrically using the Wilcoxon–Mann–Whitney U-test (Sokal, 1982). Statistical analysis and visualization were performed utilizing R Studio. Significant differences are shown as asterisks (p < 0.001 ***; < 0.01 **; < 0.05 *), otherwise as p values.

## Results

### Effects of aerosol exposure on isohydric poplar in 2021

In SEM images, the surfaces of plants grown in ambient aerosol (AA) treatment appeared different from plants grown in the aerosol-free filtered air (FA) treatment. FA leaves exhibited a rougher basic structure and stripe-like strands on the cuticular ledges. These appeared veil-like on the leaves with aerosols and depressions found on the aerosol-free leaves appeared to be filled with material, making them appear flat (Figure 2).

**Figure 2:**
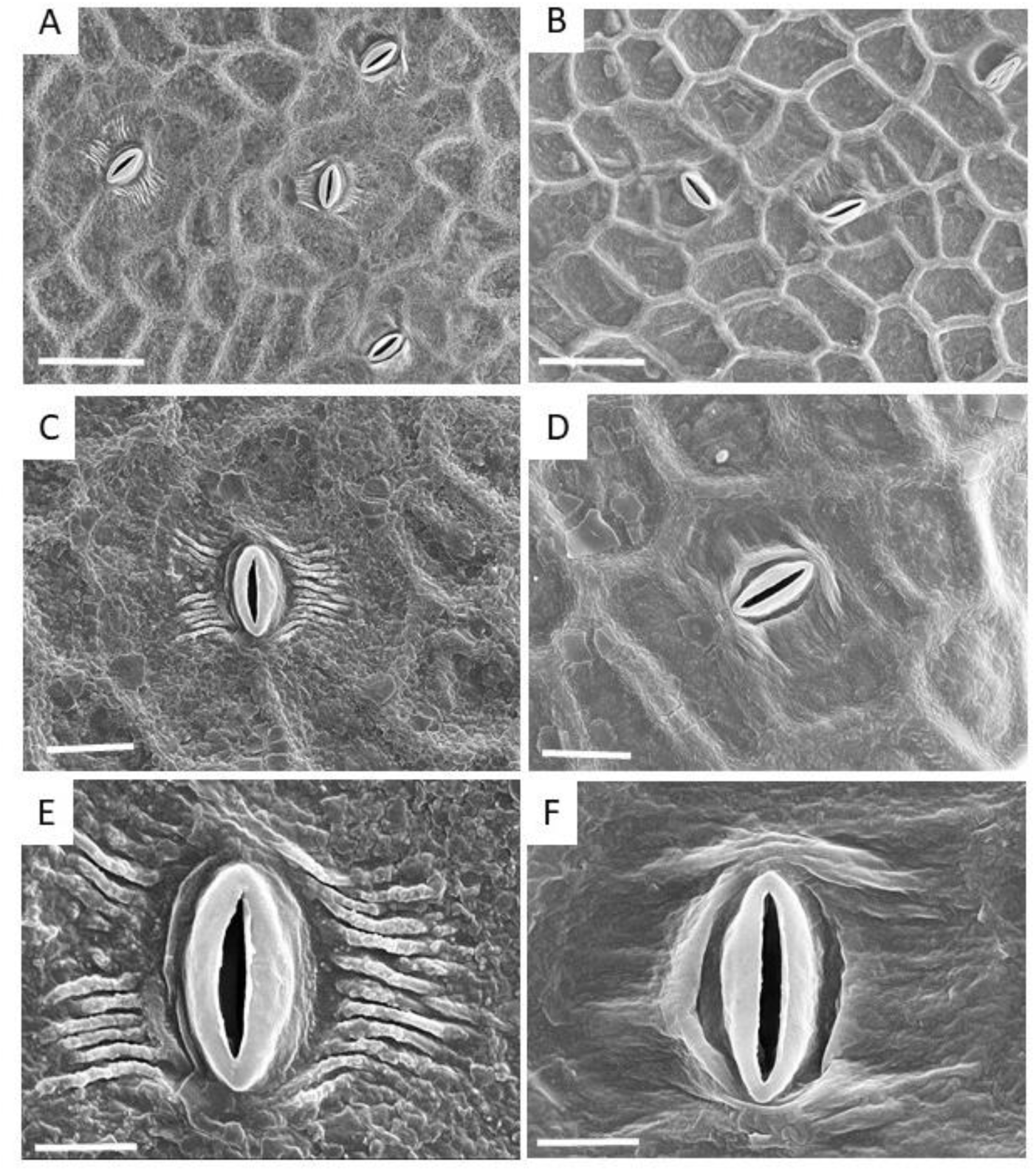
SEM images of upper surfaces of ISO leaves, showing the influence of aerosols on the surface structures on poplar leaves. Left column: aerosol-free air (FA treatment). Right column: ambient air (AA treatment). Bar length: A, B: 50 µm; C,D: 20 µm; E, F: 10 µm

The results of exploratory measurements indicated that growth of poplar clone, ISO, in the presence of ambient aerosol (AA treatment) induced several responses associated with increased water deficit (Table 1), relative to plants grown in aerosol-free filtered air (FA treatment). Osmotic adjustment, reflected in the turgor loss point, 𝜋_tlp_, and stomatal density, a developmentally determined morphological parameter, were lower in AA, while minimum leaf conductance, g_min_, was higher (Table 1). The 𝛿^13^C values, reflecting long-term restriction of stomatal conductance, g_sw_, and often a sign of water deficit, did not differ between the two growth environments.

**Table 1.** Effect of aerosol exposure on long-term indicators of water deficit in clone ISO in 2021. Values are means ± s.e., with significant differences (t-test) at p < 0.001 ***; < 0.01 **; < 0.05 *.

| Indicator (unit) | With aerosols | n | Without aerosols | n | Significance |
| --- | --- | --- | --- | --- | --- |
| $\delta^{13}C$ | $-32.7 \pm 0.14$ | 29 | $-32.8 \pm 0.13$ | 20 | $p = 0.60$ |
| $\pi_{tlp}$ (MPa) | $-2.92 \pm 0.04$ | 24 | $-2.58 \pm 0.05$ | 24 | *** |
| $g_{min}$ (mmol m<sup>-2</sup> s<sup>-1</sup>) | $2.53 \pm 0.09$ | 28 | $2.12 \pm 0.12$ | 19 | ** |
| St. Density (stomata mm<sup>-2</sup>) | $225.00 \pm 14.91$ | 10 | $265.00 \pm 11.30$ | 10 | * |

These preliminary observations, indicating that long-term aerosol exposure might degrade plant water relations in this isohydric genotype, suggested that further in-depth studies, and comparison with a closely related anisohydric genotype, would be informative.

### Effects of aerosol exposure on isohydric poplar in 2022

A more extensive set of potential indicators of long-term plant water deficit was evaluated in the same clone, ISO, in 2022 (Table 2). Confirming the preliminary studies in the previous year, values of 𝜋 _tlp_ and stomatal density were lower in the AA plants. Compared to 2021, the absolute g_min_ values were lower in 2022, increasing with aerosol by a similar 17%, but with fewer repetitions in 2022 and not significant (p = 0.28). Again values of 𝛿^13^C did not differ significantly, and neither did 𝜓_noon_.

**Table 2.**
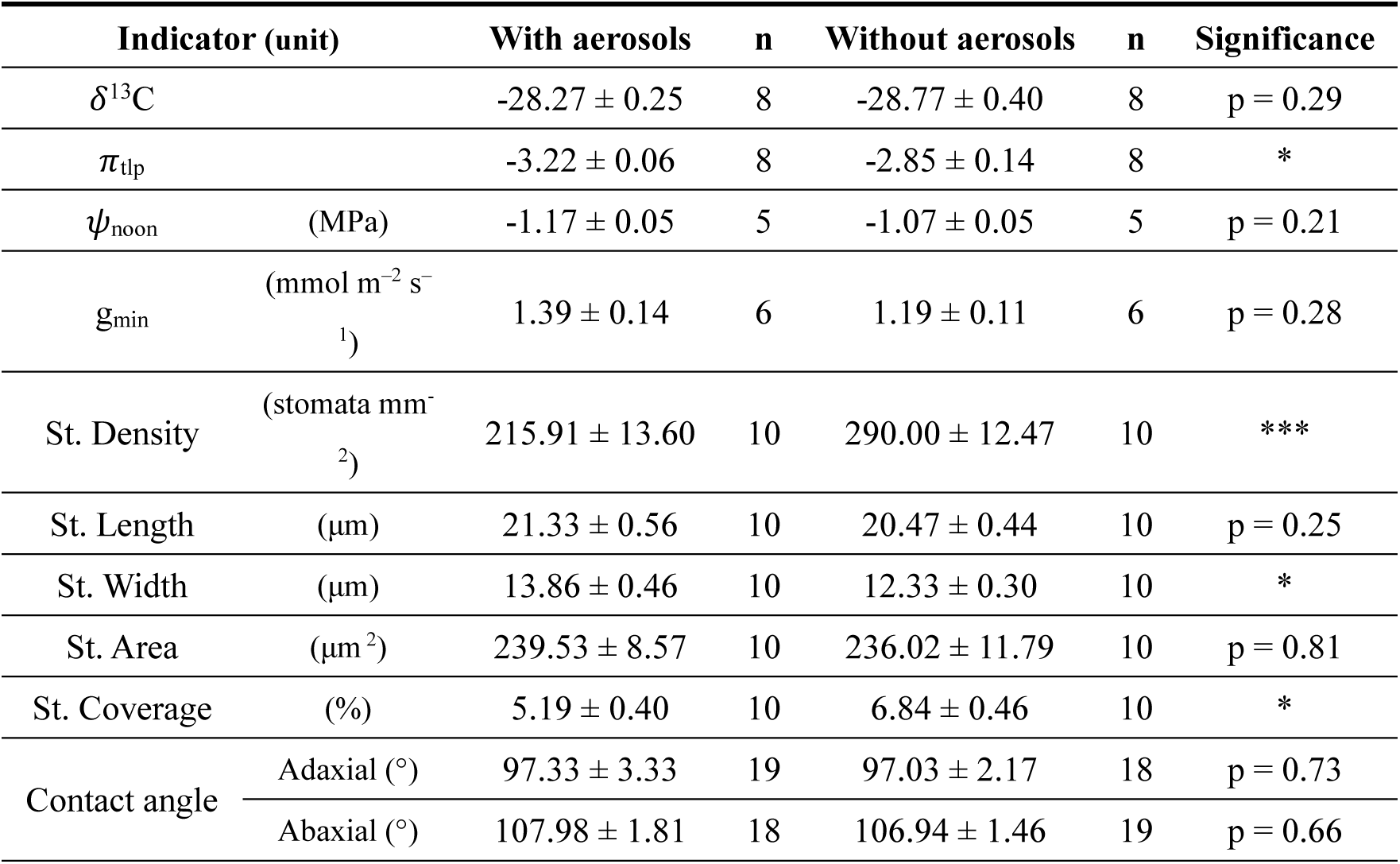
Effect of aerosol exposure on indicators of long term water deficit in ISO in 2022. Values are means ± s.e., (except the adaxial contact angle is presented as median ± s.e. of medians) with significant differences (t-test except U-test for the adaxial contact angle) at p < 0.001 ***; < 0.01 **; < 0.05 *.

### Effects of aerosol exposure on anisohydric poplar in 2022

A similar extensive set of potential indicators of long term water deficit were examined in a closely related anisohydric clone, ANI (Table 3). The 𝛿^13^C was higher (i.e., less negative), while 𝜓_noon_ was lower (i.e., more negative), in AA than FA. 𝜋_tlp_ of ANI did not differ significantly between greenhouses. The mean value of g_min_ was substantially greater in AA but did not differ significantly with few repetitions (p = 0.09; Table 3). Stomata were considerably larger and stomatal density was smaller than in ISO. Mean stomatal density was 21% lower in AA than FA, though not reaching significance (p = 0.07).

**Table 3.**
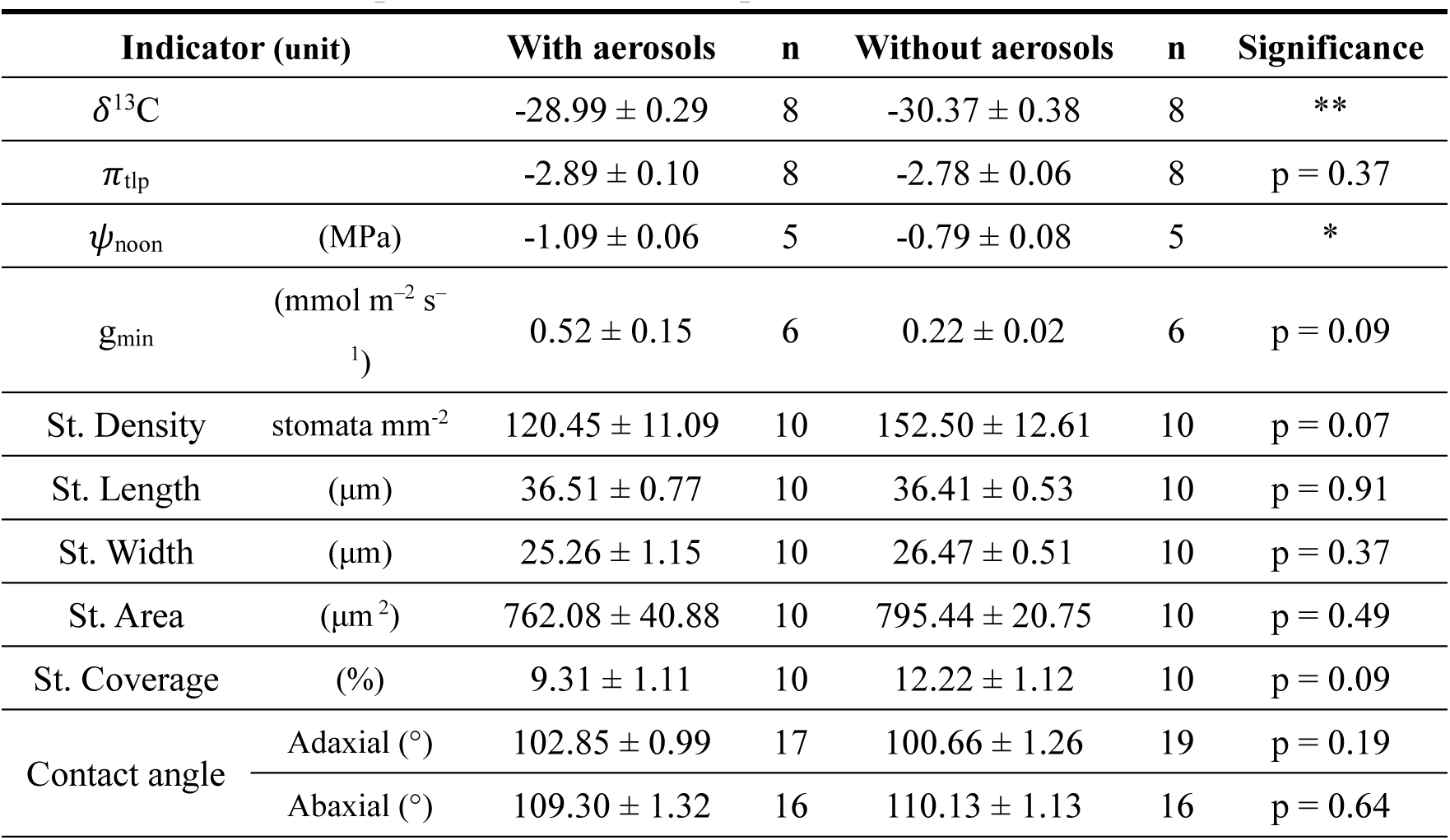
Effect of aerosols on indicators of long term water deficit in ANI in 2022. Values are means ± s.e., (except 𝛿^13^C is presented as median ± s.e. of medians) with significant differences (t-test except U-test for 𝛿^13^C) at p < 0.001 ***; < 0.01 **; < 0.05 *.

### Relative magnitude of aerosol effects on isohydric and anisohydric clones

In these data it became clear that ISO was generally less affected by aerosol than ANI. The ANI stomata were generally larger than ISO stomata and had lower density (Table 3). To visualize these differences, the values of selected parameters for each clone in AA were normalized by the mean value of the same parameter in FA (Figure 3 A-F).

**Figure 3.**
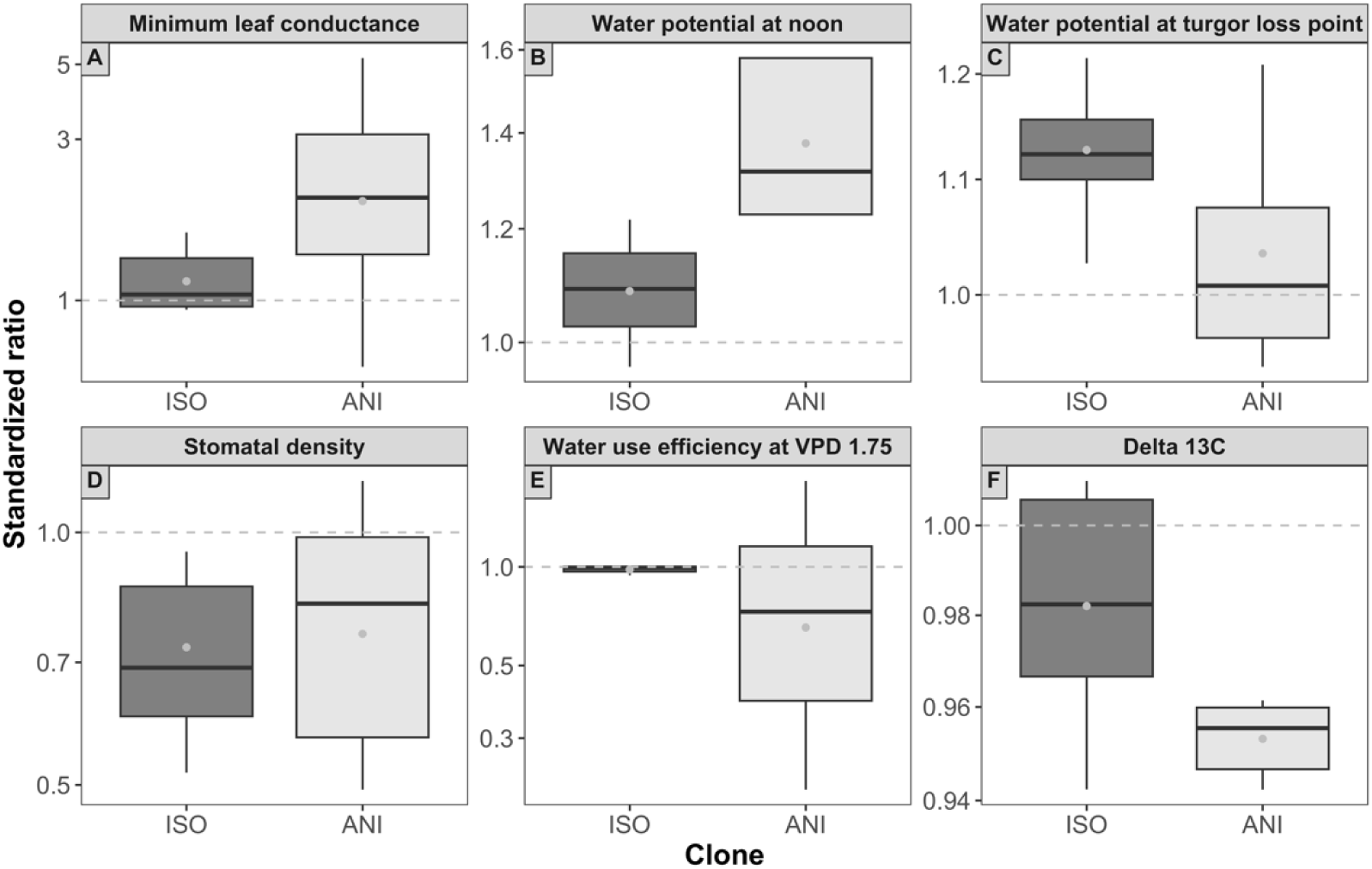
Relative magnitude of aerosol effects in two poplar clones. Ratios were obtained by normalizing individual values obtained in AA by the within-clone mean of the respective parameter in FA. The box represented the 50% of the central data, starting from the first quartile and ending in the third, with the inner line of median and the dot of mean value. The outer line on each side of the box showed the furthest data without counting outliers.

Aerosol effects on g_min_, 𝜓_noon_, WUE (observed at VPD = 1.75 kPa), and 𝛿^13^C (Figure 3 A, B, E, F) were all more pronounced (i.e. farther from the horizontal line at “ratio = 1”) in ANI than in ISO. In contrast, 𝜋_tlp_ and stomatal density exhibited greater response to aerosol in ISO than in ANI (Figure 3 C, D).

Effect of aerosol exposure on the stomatal responses to increasing VPD differed in the two clones (Figure 4A, B). The ISO clone exhibited a monotonic decline of g_sw_ of 69% in both AA and FA, but no difference due to aerosol exposure (Figure 4). The ANI clone exhibited a stronger response to aerosol, and g_sw_ at all VPD levels was reduced by AA. The decline in g_sw_ with increasing VPD resulted in a similar decrease of 29% and 32%, respectively, in g_sw_ of AA and FA. Thus in both aerosol treatments, the relative VPD response of ANI was substantially less than of ISO.

**Figure 4.**
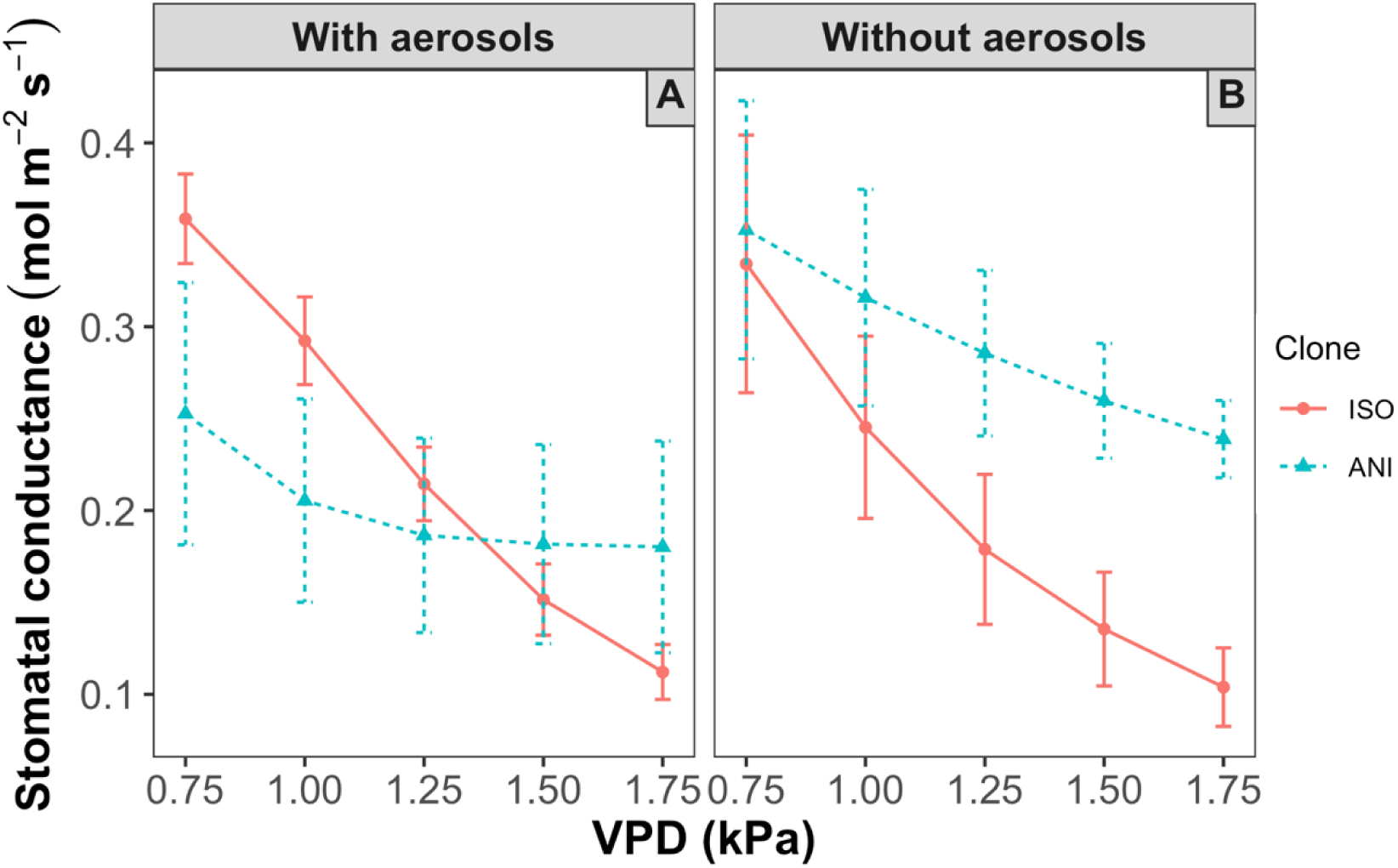
Response of stomatal conductance (g_sw_) to VPD of two poplar clones grown with and without aerosols. Data are means ± s.e. (n = 3).

Increasing evaporative demand (increasing VPD at constant T) caused the transpiration rate (E, mmol m^-2^ s^-1^; Figure 5 A, B) to increase sharply and monotonically in the anisohydric clone, ANI. In contrast, E of ISO exhibited a biphasic response with little change at moderate VPD but declining below the level of E at low VPD as VPD increased above 1.25 kPa (Figure 5A, B). This was more pronounced in AA than in FA.

**Figure 5.**
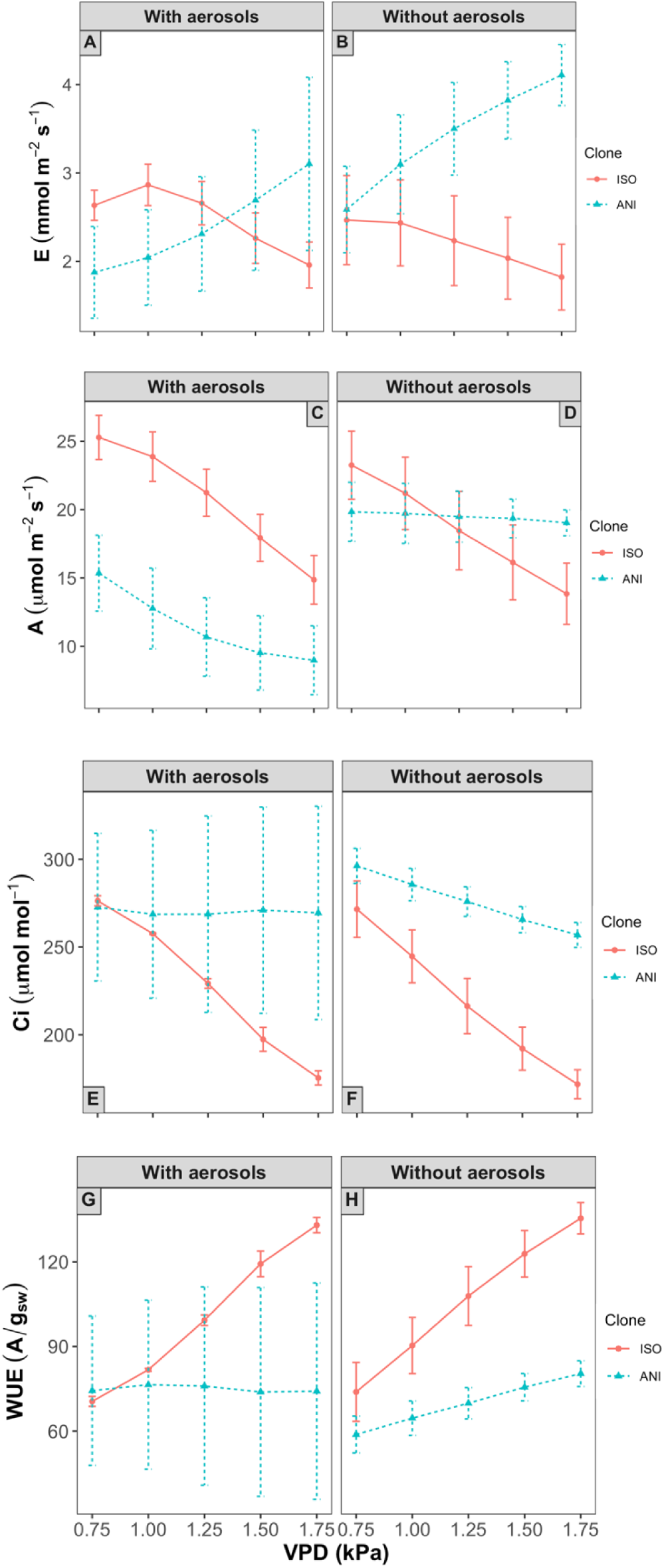
Effect of aerosol exposure on the response of E, A, Ci, and intrinsic water use efficiency, WUE, to increasing VPD of two poplar clones. Transpiration rate (A, B); Assimilation rate (C, D); Intercellular CO_2_ concentration (E, F); Intrinsic water use efficiency (G, H). Data are means ± s.e. (n = 3).

Assimilation rate (A) of ANI was reduced in AA at all VPD levels relative to FA (Figure 5C, D). In ISO, aerosol had little effect on the magnitude nor on the substantial decline of A with increasing VPD. Aerosol had a profound effect on the response of ANI to VPD (cf. Figure 5C, D). A declined by a third with increasing VPD in AA but was stable across this range of VPD in FA.

Intercellular CO_2_ concentration (C_i_) of ISO was strongly reduced by VPD but did not change in response to aerosol. Aerosol influence on C_i_ of ANI was obscured due to large variability between repetitions in AA. C_i_ of FA showed a moderate decrease with increasing VPD (Figure 5E, F).

Intrinsic water use efficiency (WUE, A/g_sw_) of ISO increased substantially with rising VPD, in both FA and AA (Figure 5G, H). In ANI, WUE response to VPD was not distinguishable in AA (Figure 5G) and exhibited a modest increase in FA

When normalized to their values at low VPD (Figure 6A, B; data from Figure 4A, B), the aerosol effect on g_sw_ of ISO was negligible, with a similar reduction at each VPD in both aerosol treatments. The final reduction was about 70%. In ANI, in contrast, the relative decline was greater at moderate VPD in the presence of aerosol, with recovery to a similar relative decline of about 30% at high VPD in both AA and FA (Figure 6B).

**Figure 6.**
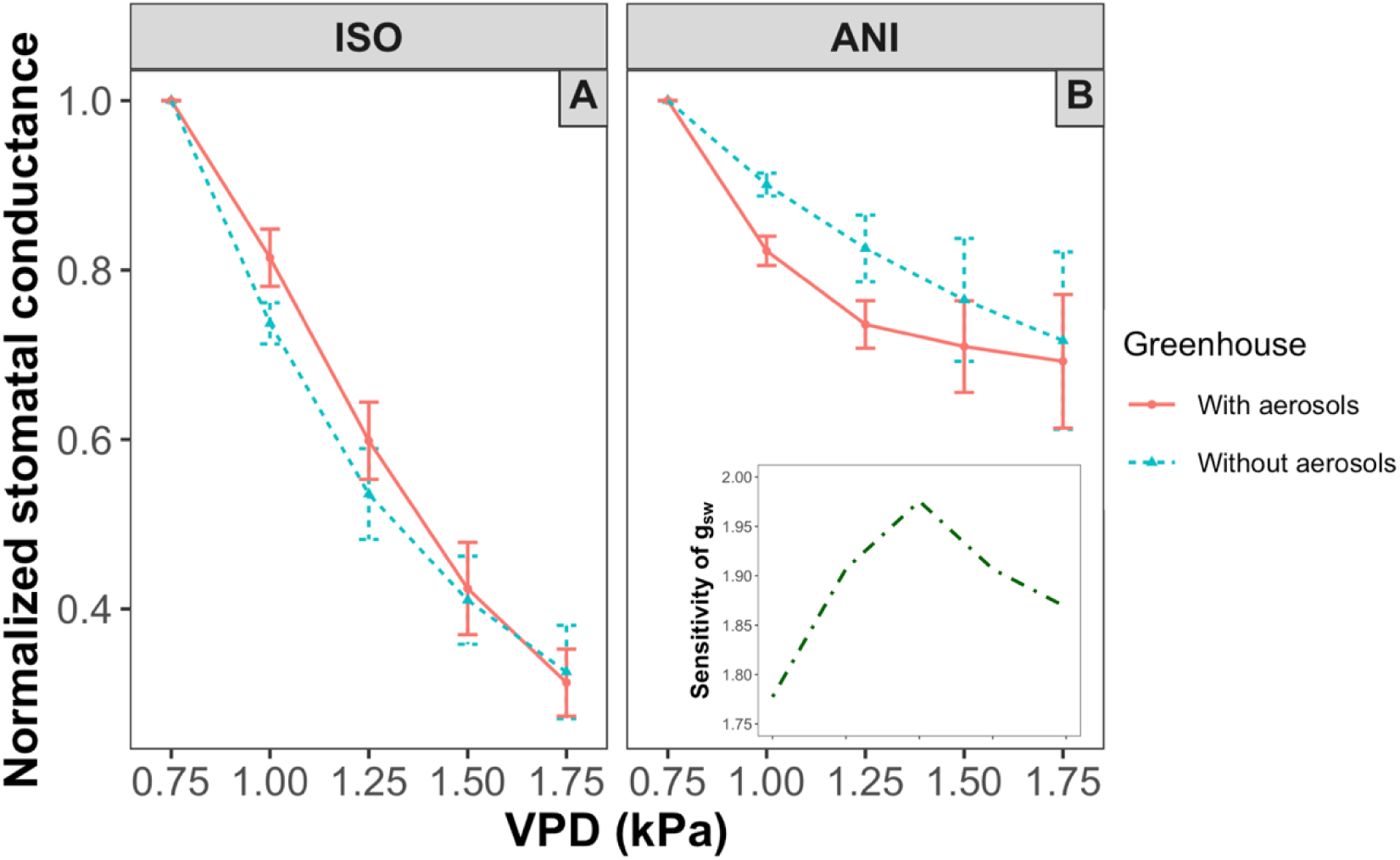
Response of g_sw_ to VPD of two poplar clones grown with and without aerosols, normalized within each curve to the value at VPD = 0.75 kPa. Data are means ± s.e. (n = 3). Each data point is the mean of individual observations obtained in AA normalized by the mean of observations of the respective parameter obtained in FA. Sensitivity of g_sw_ in ANI, estimated from g_sw_ (FA/AA, data from Figure 3) is additionally demonstrated in the side figure in B. Data are means ± s.e. (n = 3).

A biphasic nature of the sensitivity of the response of g_sw_ to aerosol with increasing VPD was observed in ANI (inset Figure 6B). In both clones, the greatest sensitivity was observed at moderate VPD.

### The Ball-Berry model

The broadly applicable Ball-Berry model of g_sw_ yields a central parameter, the B-B Index (A_n_*H_s_/C_s_), that is related to measured g_sw_ through a linear slope, g_1_, and intercept, g_0_. In ISO there were significant linear relationships between g_sw_ and the B-B Index, but these relationships were not significantly affected by aerosol exposure (Figure 7A). In ANI, however, the slope (g1) of the significant linear relationship between g_sw_ and the B-B Index ( R^2^ > 0.90) was reduced by about half in AA (p < 0.005; Figure 7B) relative to FA. The sensitivity of g_1_ was greater in ANI, similar to the directly measured WUE (cf. Figure 3E).

**Figure 7.**
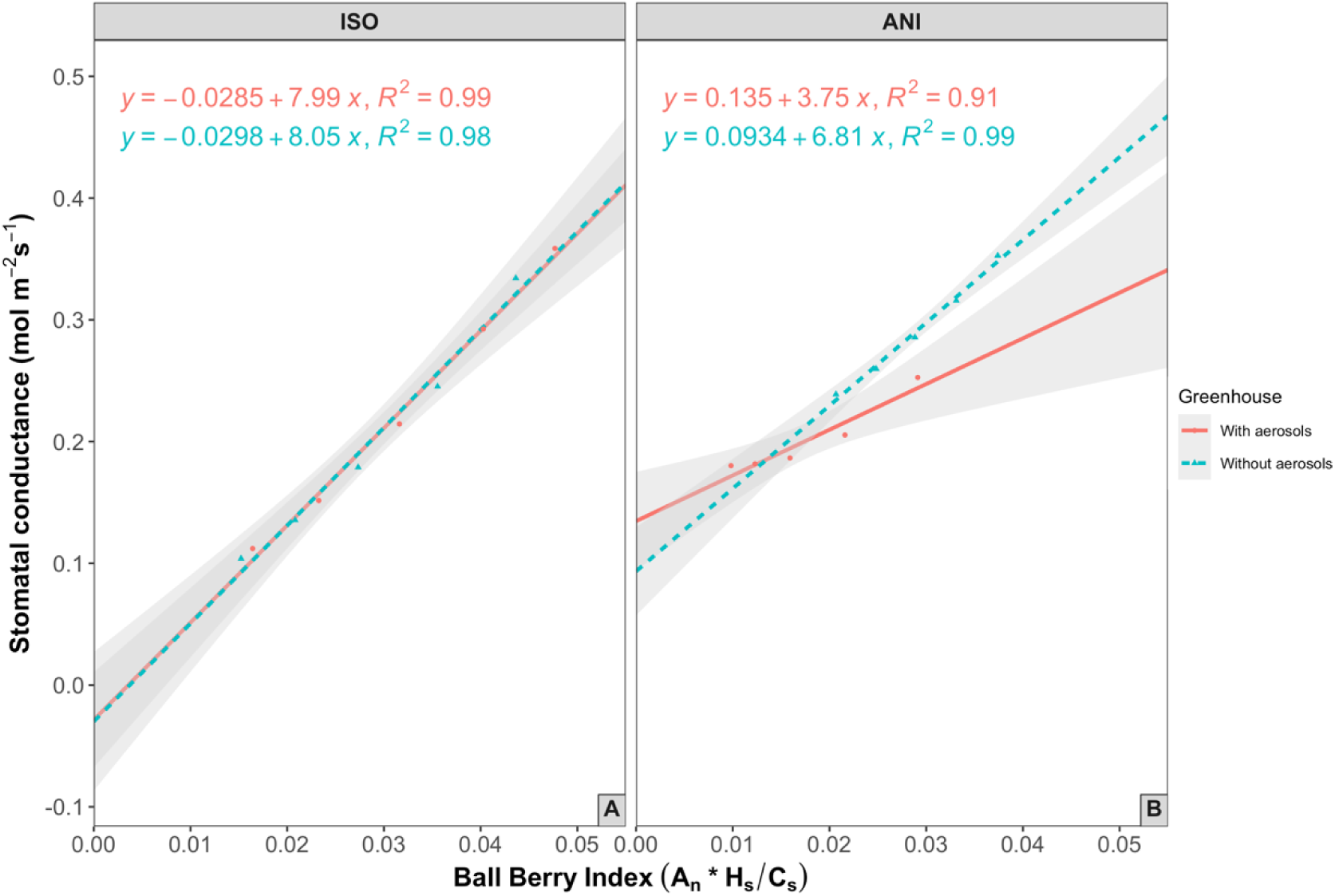
Relationship between measured stomatal conductance and the Ball-Berry Index for leaves from two poplar clones, grown with and without aerosols. The lines represent linear functions fitted to means (n = 3) of g_sw_ and the B-B Index calculated at each level of VPD. Shaded areas indicate 95% confidence intervals.

In ANI, the intercept of g_sw_ at B-B Index = 0 was somewhat greater in AA than in FA. The sensitivity of g_0_ to aerosol was greater in ANI than in ISO. This pattern in g_0_ (Fig. 7B), a measure of minimal g_sw_ when A = 0, was similar to that observed in the directly measured g_min_ (cf. Figure 3A), but the significant g_min_ difference between AA and FA plants of ISO (Table 1) was not reflected in the BB approach, which extrapolates from gas exchange data.

### Differential responses of hydroponic poplars and sunflowers

Table 4 shows the short-term responses to VPD and aerosols (PM), i.e., gas exchange and water potential. Table 5 shows the long-term responses to aerosols (Ball-Berry parameters and derived WUE; carbon isotopes, minimum leaf conductance, turgor loss point, stomatal density, TPU limitation, leaf mass per area), summarizing the results of the two poplar clones and also including data of the comparable aerosol exclusion experiment with hydroponic sunflowers (Burkhardt et al. 2023). FA sunflowers are more anisohydric than FA ANI, based on less than 1% g_sw_ decrease with VPD increasing from 0.75 kPa to 1.75 kPa (Fig 2B in Burkhardt et al., 2023) compared to ∼ 30% decrease in ANI and ∼70% in

**Table 4:** Short-term responses to vapour pressure deficit and ambient aerosols (particulate matter, PM).

| Parameter | $g_{sw}$ (ISO, ANI); $G_{sw}$ (Sunflower) | | | E | | | A | | | WUE | | | $\Psi$ |
| --- | --- | --- | --- | --- | --- | --- | --- | --- | --- | --- | --- | --- | --- |
| Factor | VPD<sub>FA</sub> | VPD<sub>AA</sub> | PM | VPD<sub>FA</sub> | VPD<sub>AA</sub> | PM | VPD<sub>FA</sub> | VPD<sub>AA</sub> | PM | VPD<sub>FA</sub> | VPD<sub>AA</sub> | PM | PM |
| ISO | - | --- | 0 | 0 | - | 0 | 0 | -- | 0 | + | +++ | 0 | 0 |
| ANI | 0 | 0 | 0 | + | 0 | 0 | 0 | 0 | -- | + | 0 | 0 | -- |
| Sunflower | -- | -- | ++ | ++ | ++ | ++ | na | na | 0 | na | na | na | 0 |

**Table 5:** Long-term responses to ambient aerosols

| Parameter | $g_0$ | $g_1$ | WUE<sub>BB</sub> | $\delta^{13}C$ | $g_{min}$ | $\pi_{TLP}$ | SD | TPU | LMA |
| --- | --- | --- | --- | --- | --- | --- | --- | --- | --- |
| ISO | 0 | 0 | 0 | 0 | ++ | -- | -- | na | na |
| ANI | 0 | --- | +++ | ++ | + | 0 | - | na | na |
| Sunflower | 0 | 0 | 0 | -- | na | -- | na | - | ++ |

ISO. The g_sw_ response to aerosols was an increase in sunflower, while no significant effects in ISO and ANI were found. Transpiration of sunflower increased more strongly than did that of ANI, both with VPD and aerosols, but differed from ANI in showing no response in water potential, B-B parameters (calculated from ACi curves) or water use efficiency. Water scarcity in sunflower in response to aerosols was noted from decreasing 𝜋_tlp_, TPU limitation, and leaf mass per area (LMA; not measured in poplars), while carbon isotopes reacted differently between ISO (0), ANI (+, less negative), and sunflower (−, more negative).

## Discussion

Both clones experienced aerosols as an additional, VPD independent, atmospheric water sink. Despite somewhat higher VPD in the FA greenhouse (Fig. 1), signs of water scarcity were consistently more apparent in AA plants. These signals were observed despite hydroponic conditions, which suggests a role for hydraulic conductivity, presumably in the leaf, which appears to be limiting (Kleaf; Brodribb & Holbrook, 2003; Sack et al., 2003; Nolan et al., 2017). This was not investigated here. FA plants experienced a ∼ 1.5 °C higher average temperature, which was not noticeable in phenological development. Because aerosol concentration was the only major factor that differed between the greenhouses, the reduced water status of AA plants can be attributed to the presence of ambient aerosols. It is assumed that optical effects due to aerosol scattering were not relevant within the greenhouse, so that the interaction between aerosols and plants occurred through aerosol deposition.

### Aerosols on the leaf surface and HAS

Aerosol deposition on poplar leaf surfaces was recognizable by comparison of SEM images from the two greenhouses. Aerosols on AA leaves appeared as flat, unstructured coverings over the pre-existing structures. Comparable structural changes are known from conifers, in which air pollution or applied particles equally caused ‘wax degradation’, whose origin is presumably also linked to covering by deliquescent aerosols (Turunen and Huttunen, 1990; Takamatsu et al., 1991; Burkhardt and Pariyar, 2014).

The coverings likely represented continuous salt crusts/solutions that developed from deliquescence of hygroscopic aerosols on the leaf surfaces, although other factors like bacterial biofilms might also be present (Morris et al., 1997). Salts deliquesce by humidification of the leaf boundary layer from transpiration and can form thin liquid films (estimated thickness 100 nm; Burkhardt, 2010) which may intrude and line the throats of stomata. This process, i.e., HAS, likely is supported by smaller size and higher density of stomata, which makes ISO more susceptible than ANI.

### HAS mediated physiological effects of aerosols

Caused by aerosols and HAS, *Vicia faba* had previously shown increased minimum leaf conductance as well as disproportionate transpiration with regard to stomatal aperture (Grantz et al., 2018). Both phenomena and several resulting consequences were also reflected in the responses of the poplar clones.

The short-term responses of the ANI clone to aerosols were

i. decreasing water potential, consistent with an anisohydric strategy (Klein, 2014; Meinzer *et al*., 2016).
ii. higher sensitivity of stomata to VPD changes: The photosynthesis of ANI plants in the FA greenhouse remained constant with increasing VPD. This changed in the AA greenhouse, where photosynthesis at 0.75 kPa VPD was 20% lower than in FA and then further decreased with increasing VPD by another 30%. Aerosol and VPD thus showed mutual enhancement.
iii. partial closure of stomata, seen by the 20-30% reductions of A and gsw, depending on VPD. Less negative δ^13^C in AA than FA also supported the decreasing stomatal aperture from aerosols.
iv. differing A and E responses. Transpiration decreased (slightly, not significantly) less than photosynthesis, leading to a 45% reduction in the g1 parameter (slope) from the Ball-Berry approach. This can probably be considered an experimental confirmation of stomatal liquid water transpiration induced by HAS. Reduced g1 also indicated increased water use efficiency (A/gsw), which is a consequence of lower stomatal aperture (cf. Chi et al.,2022; Knauer et al., 2017, Hetherington and Woodward, 2003).

The stomata of ISO closed extremely sensitively, leading to 70% reduction in gsw with the VPD increase from 0.75 to 1.75 kPa. Robust stomatal closure with increasing VPD is a defining characteristic of isohydric behaviour and may have masked small aerosol effects on g_sw_ and other parameters. However, the “feedforward response” of ISO showed an almost 30% reduction of E (p=0.061) when VPD increased from 1 to 1.75 kPa and was only observed in AA but not in FA plants. This gives experimental support for the role of aerosols in the “feedforward response”. Deposited aerosols and HAS play a role as “humidity sensor” by reacting to outside relative humidity and hydraulic transmission to the interior of the leaf. Simultaneously this introduces a functional role for the parameter Hs (relative humidity at the leaf surface) in the Ball-Berry mode. Aerosols and salts equilibrate immediately with the surrounding relative humidity and react by water absorption and desorption. This is probably the first consistent physical explanation for the scaling factor Hs in the Ball-Berry formalism, and explains its success, supporting the third hypothesis.

While in the ISO clone there was less evidence for stomatal short-term responses to aerosols than in ANI, the minimum leaf conductance g_min_ increased significantly by aerosols in 2021. Alternative mechanisms causing increased g_min_, such as increased cuticular conductance in the AA greenhouse due to the slightly lower temperature and resulting higher relative humidity (Tredenick *et al*., 2018) may be contributory. However, the g_min_ measurements of poplar clones from both aerosol environments were obtained under the same laboratory conditions over a period of several hours, equalizing cuticular conductance, and previous studies with multiple species in this same ambient exposure system support that indeed g_min_ was increased by aerosol (Burkhardt *et al*., 2001; Grantz *et al*., 2018).

The osmotic potential at turgor loss (𝜋_tlp_) and stomatal density were both significantly reduced in ISO when comparing AA with FA. Adjustment of 𝜋_tlp_ has been related to plant drought tolerance (Marechaux *et al*., 2015) and to anisohydric water use strategies under soil drought conditions (Eller *et al*., 2016; Nolan *et al*., 2017), but here was more evident for ISO than ANI. The greater osmotic adjustment in ISO could come from greater metabolic sensitivity to water deficit in the isohydric clone than in ANI, or from more sustained water deficit (McDowell, 2011; Martinez-Vilalta & Garcia-Forner, 2017). Aerosols probably caused sustained water deficit by increasing g_min_.

Reduced abaxial stomatal density is a well-established drought symptom in poplar (e.g. Himes et al., 2021; Rosso et al., 2023), which declined in both clones (by 26%; p<0.001 in ISO and by 21%; p = 0.07 in ANI) in the presence of aerosols. Stomatal density has been negatively correlated with VPD and long-term water status in poplar, where the VPD signal sensed by the basal, fully developed leaves was shown to affect the stomatal density of the upper, developing leaves (Miyazawa *et al*., 2006; Monclus *et al*., 2006). Aerosol thus acted equivalent to increasing VPD, supporting the first hypothesis.

The Ball-Berry slope (g_1_) of ISO was unaffected by aerosol, consistent with directly measured WUE and supporting its use as a predictor of WUE (Knauer *et al*., 2017; Miner & Bauerle, 2017). g0 was also not affected by aerosols, despite higher gmin. This supports that the Ball-Berry formalism can be a useful, operational tool to detect aerosol influence and HAS, but requires plant species with suitable stomatal sensitivity, like ANI.

### Plant response types

VPD increase and the presence of aerosols, each alone but also mutually reinforcing, led to consistent signs of water scarcity in each of the poplar clones and manifested in a wide range of symptoms. ANI and ISO differed in their responses. Decreasing water potential showed that ANI was anisohydric but not as strongly as sunflower, which exhibited a more pronounced increase in transpiration and conductance in response to aerosols (Burkhardt et al., 2023). The contribution of the leaf surface aerosols to transpiration is probably reflected in the difference between g_sw_ curves of AA and FA (Fig. 6), which translates to bi-phasic behaviour in ANI. The region with highest sensitivity is the VPD range where the leaf surface aerosols are most interacting with gas exchange. At low VPD, aerosols do not release water to the atmosphere due to their low saturation vapour pressure. At high VPD, air humidity is too small and aerosols are dry. At intermediate VPD, aerosols interact with water (vapour) by wicking, absorption, and desorption, particularly if there is air movement causing small local humidity fluctuations.

Together, hydroponic ISO, ANI, and sunflower showed different response types to aerosols, depending on their respective water strategy, and thus supported the second hypothesis. Atmospheric dryness may cause i) increasing transpiration in strongly anisohydric plants, like sunflower (Burkhardt et al., 2023), or ii) decreasing stomatal aperture and decreasing water potential, like ANI, or iii) very sensitive stomatal response to increasing VPD but poorly tuned to the presence of hygroscopic aerosols, like ISO.

### The risk of isohydricity

Normally, isohydric plants like ISO are thought to be more drought resistant than ANI.

Denser stomata tend to be smaller and faster at closing or opening in response to changing environmental conditions, allowing more efficient gas exchange resulting in reduced risk of hydraulic dysfunction (Durand et al., 2019; Hetherington & Woodward, 2003; Drake et al., 2013; Rosso et al., 2023). A field trial including 18 poplar hybrid clones showed a direct correlation between abaxial stomatal density and drought resistance (Himes et al., 2021; Rosso et al., 2023). Yet, isohydric trees are more severely affected by tree mortality (Breshears et al., 2005; McDowell et al., 2008) that now appears not to be a consequence of carbon starvation.

Persistently higher g_min_ from aerosols and HAS may transfer additional drought stress to the plant interior and cause metabolic responses. Accumulation is an additional problem, particularly on long-lived conifer needles. In our experiments, despite well-developed VPD sensitivity, ISO was poorly tuned to aerosol, even in the low VPD range. Ambient aerosol decreased π_TLP_, suggesting chronic stress, which these drought-avoiding plants cannot escape. These all support the fourth hypothesis. These aerosol effects should be considered in the search for reasons why isohydric trees like pinon are more damaged than anisohydric species like juniper (e.g., Breshears et al., 2005), and more broadly in models of plant water budgets.

## Acknowledgements

We acknowledge Angelika Glogau, Angelika Veits, Deborah Rupprecht, Nur Goemec from the Institute of Crop Science and Resource Conservation, and Yaron Malkowsky from Nees Institute for Biodiversity of Plants, University of Bonn, Germany, for assisting on the analytical instruments. In addition, we acknowledge Paul Gruenhofer from Department of Ecophysiology, IZMB, University of Bonn, Germany, for supporting the plant materials arrangement. This research was funded by Deutscher Akademischer Austauschdienst (DAAD, German Academic Exchange Service), grant numbers: 57440921. JB was funded by the Deutsche Forschungsgemeinschaft (DFG, German Research Foundation), grant number: 446535617. DAG was supported by the Deutsche Forschungsgemeinschaft (DFG, German Research Foundation), grant number: 446535617 and unrestricted research funds provided by the University of California at Riverside.

## Author contributions

JB and CEC developed the research design and methodologies. CEC conducted the experiment with inputs from JB. CEC and JB performed data analysis with inputs from DAG. CEC wrote the original draft. JB and DAG contributed with review and editing. All authors have read and agreed to the submitted version of the manuscript.

